# Relating genome completeness to functional predictions

**DOI:** 10.1101/2021.10.01.462806

**Authors:** Jessica Liu, Tre’Andice Williams, John A. Burns

## Abstract

Genome and transcriptome assemblies vary in their quality in terms of the connectedness of the assemblies and the amount of biological information captured. Interpreting *de novo* assemblies from new, poorly characterized, organisms in the context of complex traits can be challenging because, in the absence of a reference, it is difficult to know how much information is enough to claim the presence or absence of a trait. This study uses randomly downsampled proteome files to compare a genome completeness metric, BUSCO, to functional predictions of the complex trait of phagocytosis in known phagocytotic organisms broadly across the eukaryotic tree of life. We find that as additional proteins are added, BUSCO scores increase incrementally, while the phagocytosis prediction follows a sigmoidal curve. Generalizing our findings, we suggest a threshold of the number of BUSCOs detected above which one would expect an accurate prediction, positive or negative, of the complex trait of phagocytosis. While these findings are specific to a single trait, the methods can be extended to consider additional functional traits and predictive frameworks.

## Introduction

Whole genome and transcriptome sequencing are becoming an integral part of the biologist’s toolkit. Sequencing of nucleic acids provides a window into the functional and evolutionary processes of an organism^1^. Every sequencing effort is not equal, though. Technical and conditional variation can affect the quality and completeness of a sequencing data set, limiting the amount of information that can be extracted and the depth of inference that can be made^2^. This is particularly true for *de novo* sequencing and assembly of whole genomes or transcriptomes of new species where no reference genome or transcriptome is available.

Evaluating the quality and completeness of a *de novo* sequencing effort involves assembly metrics like read depth, assembly length, and functional content of the final assembly^3^. To get a sense for how well an assembly captures the protein coding space of a new organism, comparative tools are used that compare the new assembly to well annotated, existing assemblies. Tools such as CEGMA^4^ and BUSCO^5^ operate on a principle of counting the number of expected core genes, defined as genes shared across taxonomic groups, that can be found in a new assembly. The counts assess the “completeness” of an assembly with respect to a standard data set.

Count-based metrics provide an overall assessment of the proportion of expected genes captured, but they lack direct relevance to functional and phenotypic properties of an organism. Researchers often want to know whether an organism has the genetic capacity to complete a particular function, and whether their *de novo* assembly contains sufficient information to confidently assess absence of a trait. For traits that can be predicted by the presence of one or two genes, genome completeness and the probability of detection of such traits should scale linearly, but for traits that are predicted by complex combinations of hundreds of genes, the relationship between genome completeness and the amount of information needed to assign absence of a trait is more nuanced^6,7^.

Here we address the question of how a genome completeness metric co-varies with a functional trait prediction for a set of diverse eukaryotic proteomes. For genome completeness, we consider BUSCO scores, and for a functional trait we use phagocytosis, a complex process that is predicted by species-specific combinations of hundreds of genes^8^. We anticipate that our analyses will help researchers better understand whether their de novo assemblies contain sufficient information to make presence/absence calls regarding complex processes like phagocytosis.

## Materials and Methods

### Proteome data

Whole proteome data were obtained from the EukProt collection^9^. The 36 species (Table S1) were selected as a diverse collection of organisms that spanned the eukaryotic tree of life (Figure 1) and are known to use a phagocytotic mode of nutrient acquisition or to carry specialized phagocytotic cells.

**Figure 1.**
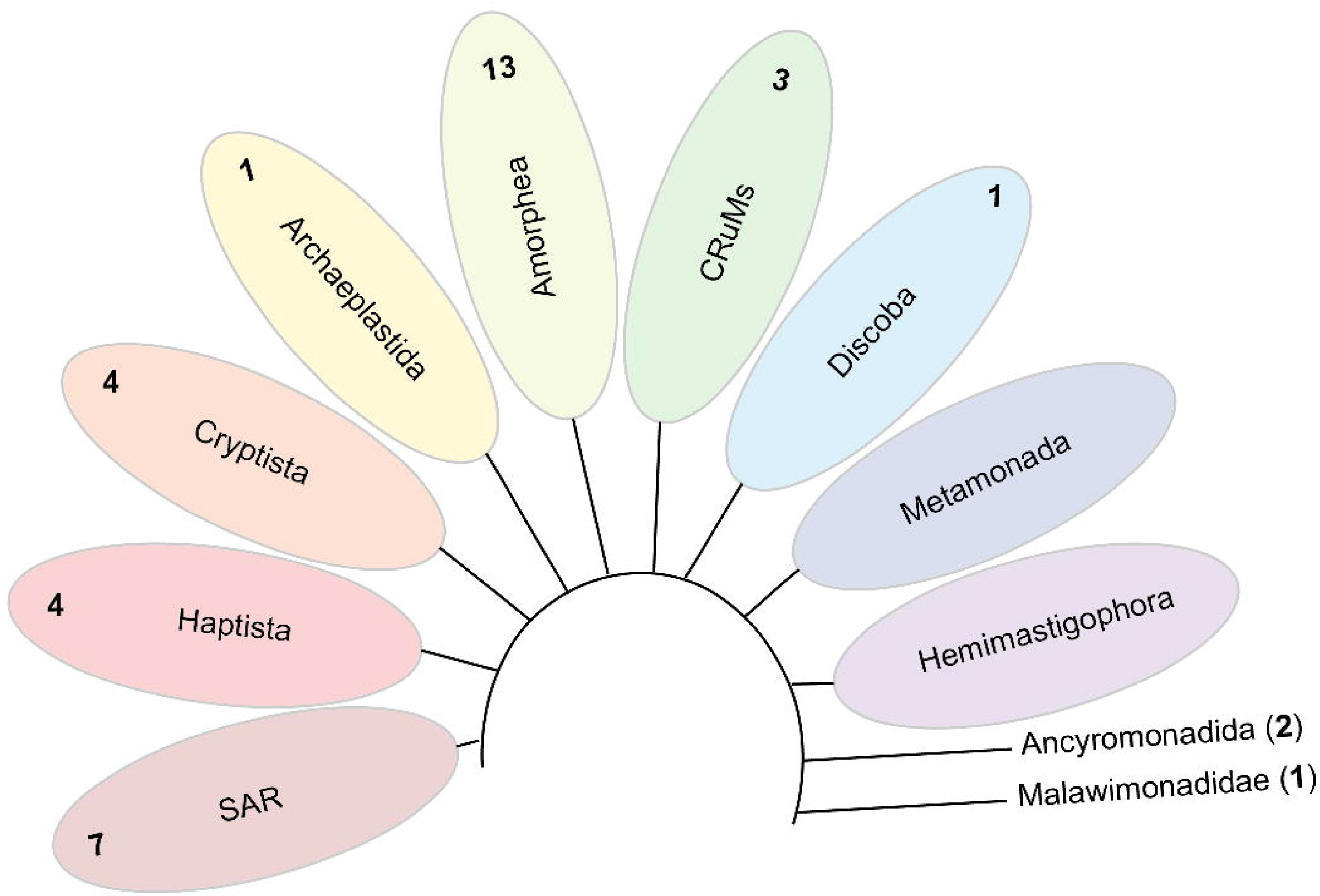
Representation of proteome data from major eukaryotic supergroups and orphan taxa. The numbers in bold are the number of taxa from each group used in the combined analyses.

### Subsampling

Proteome files were subsampled by taking a random selection of fasta entries equivalent to a percentage of the total proteins in a given proteome using a custom bash script (see: GenomePercPredict^10^). Twenty replicate subsamples were taken for each increment of proteome completeness from 5% through 95% complete at 5% intervals.

### Phagocytosis predictions

For each complete proteome, peptides were searched against the 14,095 HMMs of the “predictTrophicMode” model^11^ using HMMER (v3.3.2)^12^. Best hit HMMs were assigned to each peptide where a match had a full sequence e-value≤ 10–5 and a best domain e-value hit≤10–4, as in Burns 2018^8^. A map of protein ID and best hit HMM was used to define sets of HMMs present in proteome subsets. HMMs mapping to peptide file subsets were used as input for the predictTrophicMode algorithm to define a phagocytosis prediction probability score for each peptide subset.

### BUSCO completeness

The BUSCO algorithm (v5.2.2)^13^ was run against each peptide subset using the eukaryota_odb10.2019-11-20 BUSCO set for all species and the number of missing BUSCOs was compiled from the BUSCO output. The number of BUSCOs detected for each peptide subset was calculated as total BUSCOs - missing BUSCOs.

### Statistics

A phagocytosis prediction score and number of BUSCOs detected was compiled for each peptide subset from each species. For individual species, the number of BUSCOs detected and phagocytosis prediction scores were plotted against proteome completeness. To summarize the breadth of patterns observed and generalize to novel genomes/transcriptomes/proteomes, we plotted all subsets of all species phagocytosis prediction scores against the number of BUSCOs detected. From this relation, we considered the empirical probability that at a given number of BUSCOs detected a genome of a true phagocyte would have a phagocytosis completeness score >0.5, indicative of a positive phagocytosis prediction, calculated as (#predictions>0.5)/(total predictions) per #BUSCOs detected. We found that the distribution of probabilities has a best fit to a 3-parameter, type-2 Weibull distribution using R-package drc v3.0.1^14^. Model fitting and plotting was performed in R v4.1.0^15^ with ggplot2 v3.3.5^16^.

## Results

By subsampling the protein space of 36 diverse phagocytotic organisms (Figure 1), we show the relationship between BUSCO scores, as a measure of genome completeness, and functional prediction probability scores for the complex process of phagocytosis (Figure 2A and B). Using the subsample data, we calculated the empirical probability that any phagocytotic organism would have a phagocytosis probability score greater than 0.5, indicating the genetic capacity for phagocytosis, given the number of BUSCOs detected (Figure 2C). We see that when 150 or more eukaryote BUSCOs were detected (complete or fragmented, but not missing), there was a high probability (98.2% ± 6.6%) that the genome contained enough information to make an accurate functional call regarding phagocytosis.

**Figure 2.**
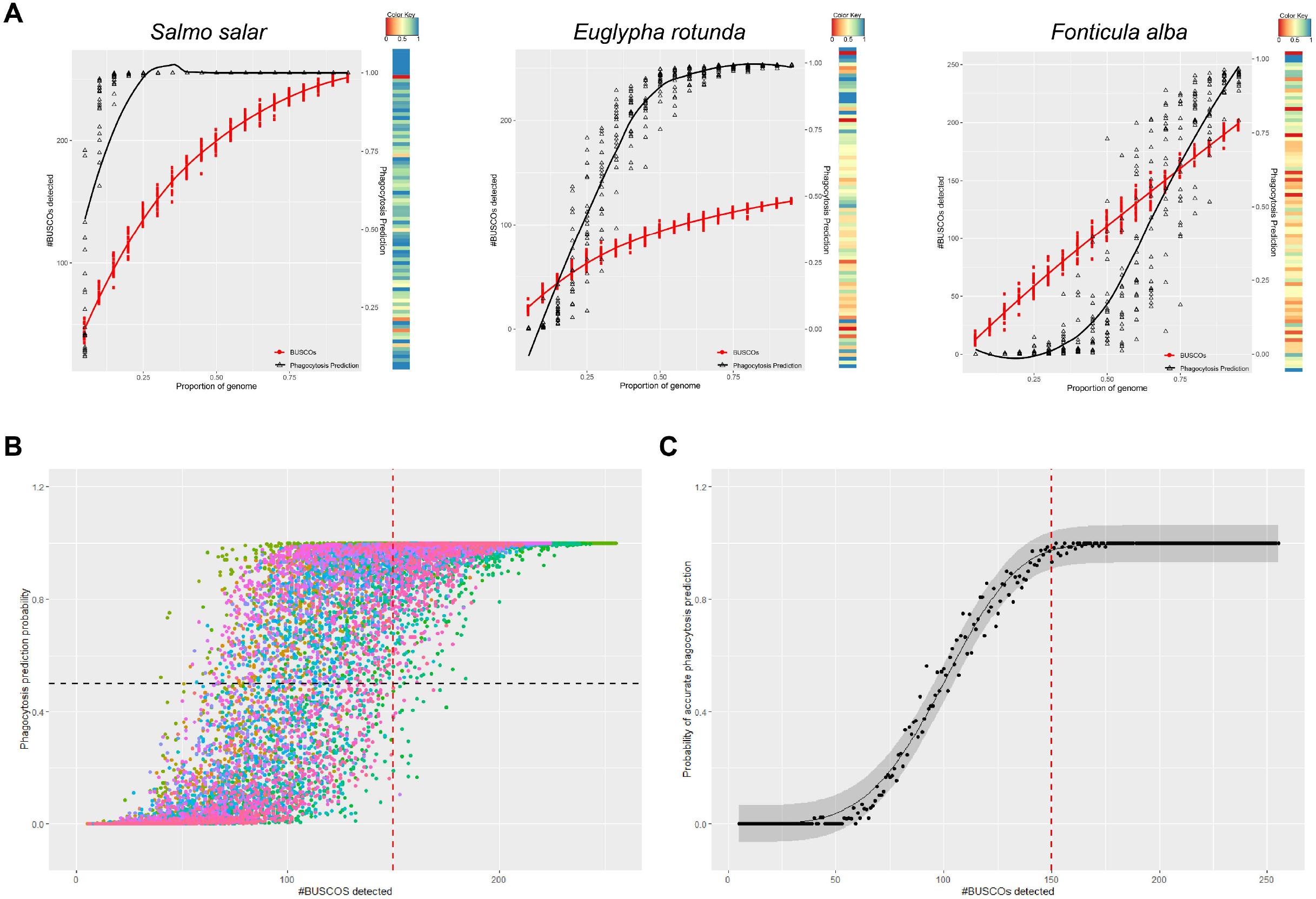
The relationship between proteome completeness, BUSCO scores, and functional predictions. **A.** Number of BUSCOs detected and phagocytosis probability vs. proportion of proteins analyzed for three select taxa. The color bar to the right of each plot shows the GO category score pattern for the complete proteome. **B.** Phagocytosis prediction probability vs. number of BUSCOs detected for all subsamples of all 36 proteomes. The black dashed horizontal line indicates a probability of 50%, above which an organism would be considered capable of phagocytosis. The red dashed vertical line indicates the threshold of 150 BUSCOs detected, above which most true phagocytes have a positive prediction for phagocytosis. **C.** The probability of a positive phagocytosis prediction (>0.5) vs. the number of BUSCOs detected. The data fits a Weibull distribution (black fit line, with a 95% confidence interval). The red dashed vertical line indicates the 150 BUSCO threshold.

## Discussion

Our results show that genome completeness metrics increase incrementally as additional genetic information is added to a proteome, while prediction of phagocytic capacity follows a sigmoidal curve. At low levels of proteome completeness, there is not enough information to make a confident functional call. Functional prediction scores begin to rise rapidly once sufficient genetic information is available, and can plateau at a point where additional proteins no longer influence the prediction. The functional predictions rely on how many genes are found, which specific genes they are, and how those genes are distributed among the functional categories that make up the model. We believe this is why some predictions can be confidently made at low proteome completeness, such as the Atlantic Salmon (*Salmo salaŕ*) (Figure 2A), which reaches an average phagocytosis prediction probability of 88% ± 9.7% with just 10% of the proteome sampled (with only 75 ± 5 out of 255 BUSCOs detected). With any 10%of the protein coding genes sampled, we would accurately predict that a salmon has cells capable of phagocytosis. For organisms that carry fewer of the genes predictive of phagocytosis, like the nuclearid *Fonticula alba* (Figure 2C), a larger proportion of the protein coding genes needs to be sampled to cross the 50% prediction probability threshold. For F. alba, on average 70% of the proteome needs to be sampled to identify the organism as a phagocyte.

A general rule emerges from this pan-eukaryote analysis that if a researcher has detected 150 BUSCOs or more (less than 105 missing BUSCOs in the BUSCO output file) in their data when considering the 255 BUSCO eukaryotic gene set, there should be enough information available to make an accurate prediction regarding phagocytotic capacity. If fewer than 150 BUSCOs are detected, the probability of collecting enough information to make an accurate functional prediction with regards to phagocytosis decreases. In such cases a positive prediction always means that an organism is predicted to be capable of phagocytosis, while a negative prediction could indicate that additional sequencing is warranted. Conversely, with over 150 BUSCOs detected, additional sequencing may not provide additional functional information with respect to phagocytosis.

For researchers working on novel genomes and transcriptomes, these analyses complement oft-used genome completeness metrics^4,5,13^ by demonstrating how functional predictions vary with genome completeness across a broad swath of eukaryotic diversity and by providing a threshold below which a researcher might determine that additional sequencing is needed. The subsampling analysis used here is focused on linking predictions of phagocytotic capability to genome completeness, but it may also be useful for, and extended to, additional functional predictions.

## Data availability

All proteome files were downloaded from and are available as part of the EukProt collection^9^. Computer code and data files used to subsample proteome files and plot and model data is available on github at https://github.com/burnsajohn/GenomePercPredict.

## Author Contributions

JL, TW, and JB designed the study and completed analyses. JB wrote the manuscript.

## Acknowledgements

The authors thank Dr. David Fields and Roxana Branch for their guidance during the virtual NSF REU program in the summer of 2020, during which this work was completed.

---

**Table S1.** 36 species used in this study.

